# Curated BLAST for Genomes

**DOI:** 10.1101/533430

**Authors:** Morgan N. Price, Adam P. Arkin

**Affiliations:** Lawrence Berkeley National Laboratory

## Abstract

“Curated BLAST for Genomes” finds candidate genes for a process or an enzymatic activity within a genome of interest. In contrast to annotation tools, which usually predict a single activity for each protein, Curated BLAST asks if any of the proteins in the genome are similar to characterized proteins that are relevant. Given a query such as an enzyme’s name or an EC number, Curated BLAST searches the curated descriptions of over 100,000 characterized proteins, and it compares the relevant characterized proteins to the predicted proteins in the genome of interest. In case of errors in the gene models, Curated BLAST also searches the six-frame translation of the genome. Curated BLAST is available at http://papers.genomics.lbl.gov/curated.

## Introduction

Given the genome sequence for an organism of interest, we often want to know whether or not it encodes a certain capability and which proteins might be involved. To support this, many genomics web sites support searching for proteins whose annotations match a text query. However, annotation tools will usually provide one predicted function for each protein, and these predictions are often incorrect (i.e., (Schnoes et al. 2009)). So searching through annotations may not be the best way to find proteins that are involved in a process.

Instead, we propose that given a text query, we can identify experimentally-characterized proteins (usually from other organisms) that are relevant. Then, we can search in the genome of interest for proteins that are similar to these characterized proteins. For enzymes, this approach obviates the need to predict the substrate specificity (which is often not possible). Instead, we identify candidates that are similar to characterized proteins that have activities of interest.

We implemented this approach in a tool called “Curated BLAST for Genomes.” It relies on a collection of over 100,000 characterized proteins, and it usually takes just a few seconds per query.

## Results

### Finding a missing annotation

For example, consider searching for “perchlorate” in *Azospira oryzae* PS (Figure 1). It takes a few seconds for Curated BLAST to identify three proteins in the genome that are over 80% identical, over their full length, to the three subunits of a putative perchlorate reductase. The putative perchlorate reductase is from a related species (*Azospira oryzae* used to be named *Dechlorosoma suillum*), was identified by genetic approaches (Bender et al. 2005), and is curated in Swiss-Prot (The UniProt Consortium 2017). Two of the proteins from the PS strain have actually been demonstrated to reduce perchlorate (PcrAB; (Youngblut et al. 2016)), but this is not reflected in any of the databases. It might still seem easy to annotate the proteins in the PS strain, given that they are so similar to proteins in Swiss-Prot, but as of November 2018, neither RefSeq (Tatusova et al. 2015) nor RAST (Overbeek et al. 2014) nor KEGG (Kanehisa et al. 2016) annotate any of these proteins as perchlorate reductase. In RAST and KEGG, these proteins are misannotated as a nitrate reductase. Perchlorate reductase illustrates how it can be easier to find a protein with Curated BLAST than by using gene annotations.

**Figure 1:**
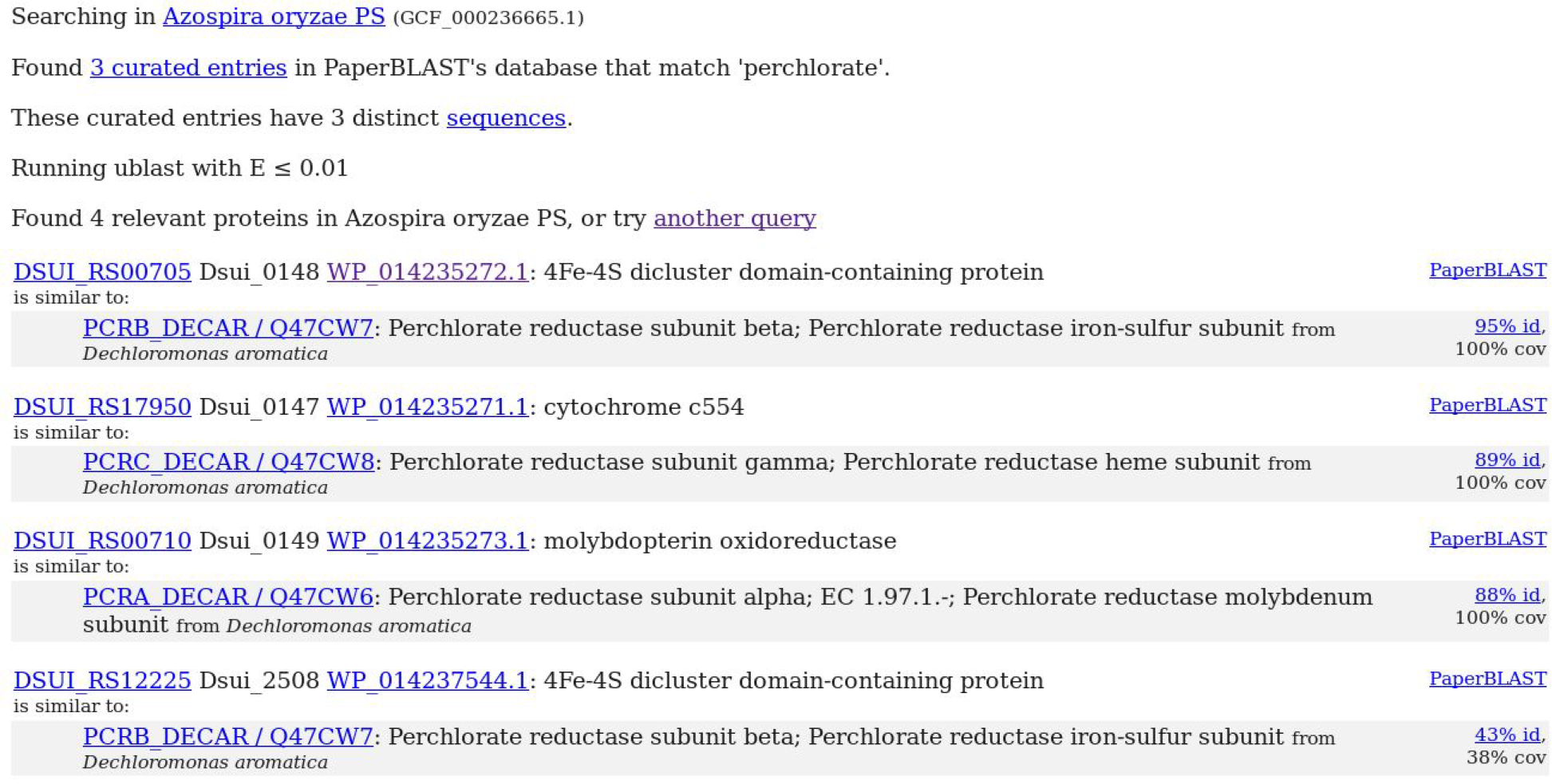
Curated BLAST’s results for “perchlorate reductase” in *Azospira oryzae* PS. “id” is short for %identity of the amino acid sequence, and “cov” is short for the %coverage of the characterized protein by the query.

Curated BLAST also found another protein in the genome of *A. oryzae*, Dsui_2508, that has some similarity to perchlorate reductase. The alignment covers less than 50% of the perchlorate reductase subunit (“38% cov”), and the similarity is modest (“43% id.”), which suggests that Dsui_2508 might have a different function. To identify other potential functions for the proteins that are returned, the search results include a link to PaperBLAST (Price and Arkin 2017), which finds papers and curated entries about homologs of a protein of interest. Clicking on the PaperBLAST link shows that Dsui_2508 is 57% identical, over 85% of its length, to a subunit of a tetrathionate reductase, so Dsui_2508 is not likely to be involved in perchlorate reduction.

### Finding candidates even when the gene models are incorrect

Another challenge in finding proteins in a genome is that the protein of interest may be missing from the list of predicted proteins. Proteomics studies often identify proteins that were missed in the computational gene annotation (i.e., (Venter et al. 2011)). Insertion or deletion errors within the sequence of a protein-coding gene can also prevent the protein from being identified. This may be a common problem with long-read single-molecule sequencing: many of the resulting genomes have a suspiciously large number of genes that are disrupted by frameshifts (Mick Watson, blog post). One way to rule out errors in gene models is to search against the six-frame translation of the genome, but six-frame searches are slow and cumbersome.

To make it easy to check for “missing” proteins, Curated BLAST searches against the six-frame translation of the genome. Because six-frame search takes several times longer than searching against the predicted proteins, Curated BLAST first searches against the predicted proteins and shows those results. It can then search against the six-frame translation while the user is inspecting the hits to the predicted proteins. Curated BLAST reports any hits to the six frame translation that were not expected given the predicted proteins. For example, we recently found that the histidinol dehydrogenase of *Azospirillum brasilense* Sp245 was not annotated (see NCBI assembly ASM23736v1) because of a frameshift error in the genome sequence (Price, Zane, et al. 2018). As shown in Figure 2, searching *A. brasilense* for “histidinol dehydrogenase” finds two nearby reading frames that are similar to characterized histidinol dehydrogenases. The two reading frames cover the N-and C-terminal portions of the protein (this can be seen by hovering on “62% id.” or “59% id.”).

**Figure 2:**
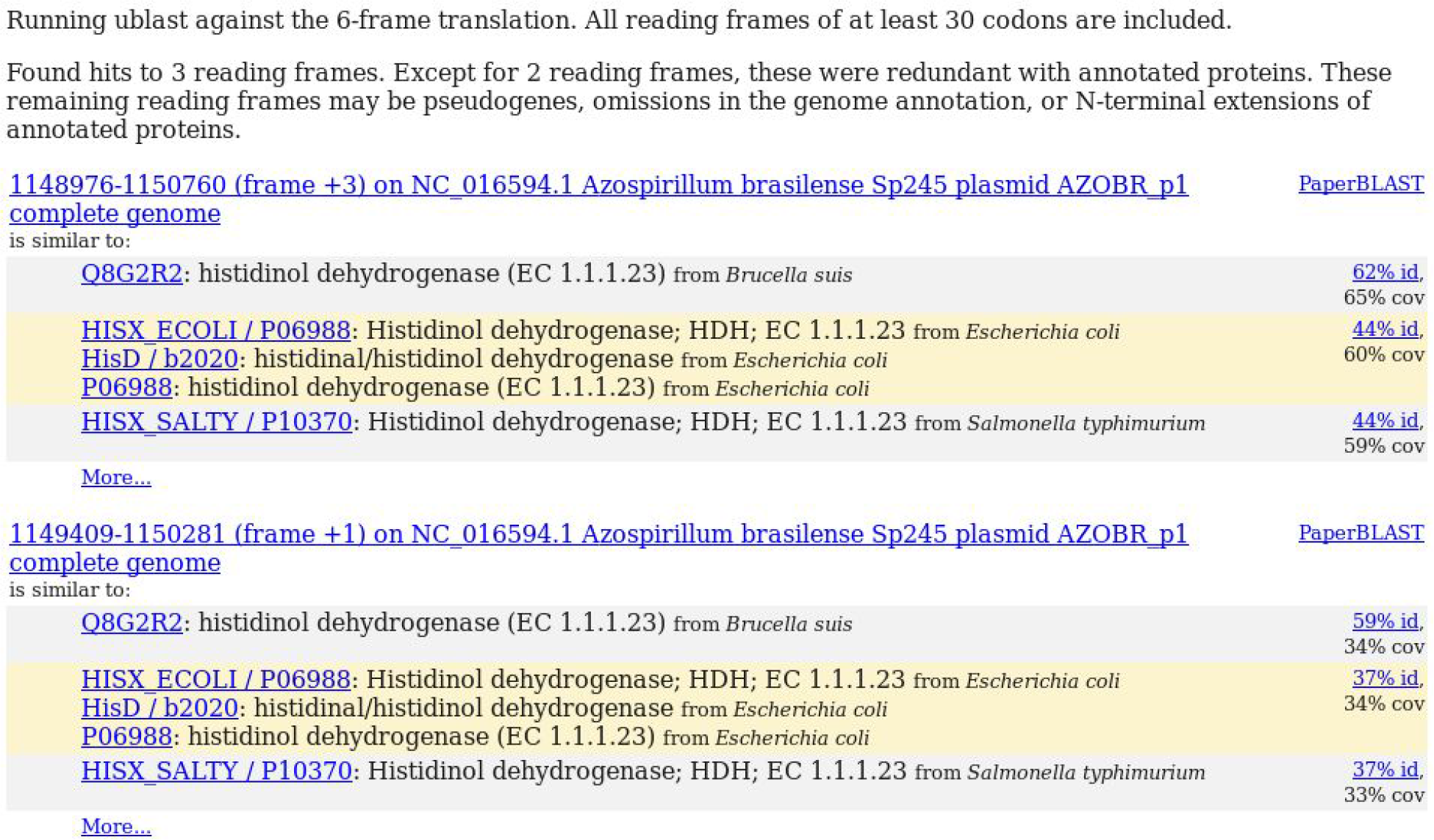
Six-frame search results for “histidinol dehydrogenase” against *Azospirillum brasilense* Sp245.

We developed six-frame search with bacterial and archaeal genomes in mind, and it does not take splicing into account. Nevertheless, it may be useful for smaller eukaryotic genomes. Six-frame search is available for genomes of up to 30 MB.

### Integration with genome databases

Curated BLAST works with genomes from NCBI’s assembly database (Sayers et al. 2018), from JGI’s Integrated Microbial Genomes (Chen et al. 2018), from MicrobesOnline (Dehal et al. 2010), or from the Fitness Browser (Price, Wetmore, et al. 2018). Curated BLAST also works with UniProt proteomes (The UniProt Consortium 2017), although six-frame search is not available for proteomes. For each of these sources, the user can search for the genome of interest by genus name, species name, and/or strain name. Alternatively, the user can upload protein or nucleotide sequences in FASTA format.

### Sources of characterized proteins

Curated BLAST relies on eight databases for curated descriptions of the functions of characterized proteins:

- BRENDA, a database of enzymes (Placzek et al. 2017).
- CAZy, a database of carbohydrate-active enzymes (Lombard et al. 2014). Only the experimentally characterized proteins are included.
- CharProtDB, a database of characterized proteins (Madupu et al. 2012).
- EcoCyc, a database of genes in *Escherichia coli* K-12 (Keseler et al. 2005).
- MetaCyc, a database of metabolism and enzymes (Caspi et al. 2010).
- REBASE, a database of DNA restriction and modification enzymes (Roberts et al. 2015).
- Swiss-Prot, the manually curated section of UniProt (The UniProt Consortium 2017). Only proteins with experimental evidence as to their function are included.
- The Fitness Browser, a database of genome-wide mutant fitness data and of proteins whose functions were identified from mutant phenotypes and comparative genomics (Price, Wetmore, et al. 2018).

Some of the proteins in CharProtDB, EcoCyC, and Swiss-Prot are not actually characterized and have vague annotations, but these vague annotations are unlikely to match a query and so they rarely affect the results. Overall, Curated BLAST’s database contains 115,874 different protein sequences, with most of these originating from Swiss-Prot (Table 1).

**Table 1:**
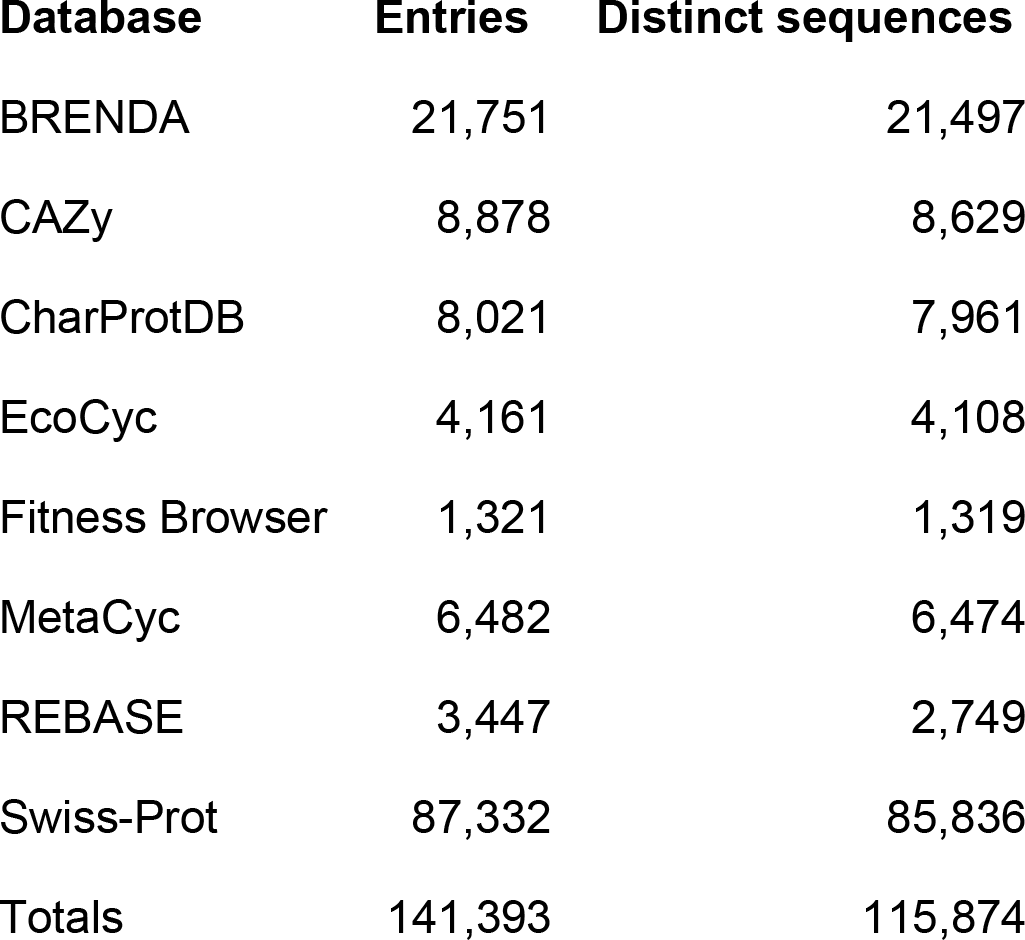
Sources of characterized proteins.

| **Database** | **Entries** | **Distinct sequences** |
| --- | --- | --- |
| BRENDA | 21,751 | 21,497 |
| CAZy | 8,878 | 8,629 |
| CharProtDB | 8,021 | 7,961 |
| EcoCyc | 4,161 | 4,108 |
| Fitness Browser | 1,321 | 1,319 |
| MetaCyc | 6,482 | 6,474 |
| REBASE | 3,447 | 2,749 |
| Swiss-Prot | 87,332 | 85,836 |
| Totals | 141,393 | 115,874 |

## Discussion

The most important limitation of Curated BLAST is the underlying database of characterized proteins. Although this database contains over 100,000 different characterized protein sequences, there are many more characterized proteins that have not been curated. Most of the microbial proteins that have been characterized in the last few years are probably missing (Price and Arkin 2017). As natural language processing technology improves, it may become feasible to extract protein functions from the full text of papers along with sequence identifiers.

Another limitation of Curated BLAST is that a text query might not include all of the characterized proteins of interest. Except for enzymes, there is no shared nomenclature for protein functions across the curated databases. For enzymes, Curated BLAST works well with enzyme commission (EC) numbers as the queries if the “word search” option is selected. (Our database links over 45,000 different protein sequences to over 5,000 fully specific four-level EC numbers.) Even so, EC numbers might be overly specific if several similar reactions are of interest. For instance, oxidizing the same substrate with different electron acceptors corresponds to different EC numbers. Also, EC numbers can change, and some of the curated entries contain out-of-date EC numbers. The results page includes a link to the list of curated genes that match the query, which you can use to see if Curated BLAST is considering all of the activities that you expect.

Another limitation is that the activity of interest may be encoded by a protein that is not related to any characterized protein that has the activity, even though the protein is similar to characterized proteins with similar activities. In this situation, searching for relevant protein families may be more effective than searching for the specific activity.

## Materials and Methods

### Software

Curated BLAST for genomes is implemented in CGI (common gateway interface) scripts and perl. It uses the same SQLite3 relational database as PaperBLAST (Price and Arkin 2017). Ublast (version 10.0) is used to compare curated sequences that match the query to the predicted proteome, with a maximum e-value of 0.01 (Edgar 2010). For six-frame translation search, all stretches of 30 amino acids or more without a stop codon are analyzed.

### Sources of characterized proteins

We previously described how EcoCyc and the characterized subset of Swiss-Prot were incorporated into the PaperBLAST database (Price and Arkin 2017). For this study, we used EcoCyc 22.5 and we downloaded Swiss-Prot on October 29, 2018. We also incorporated curated gene descriptions from BRENDA, CAZy, CharProtDB, MetaCyc, REBASE, and the Fitness Browser.

BRENDA was downloaded on January 11, 2018. Only entries that contain a UniProt identifier and at least one publication were retained. The corresponding protein sequences were obtained from UniProt. Protein sequences that were fragments (according to UniProt’s fasta files) were excluded.

CAZy was obtained in FASTA format and as a table of EC numbers (link) via dbCAN (Yin et al. 2012) on November 15, 2017. Only entries that are linked to EC numbers, which should be experimentally characterized, were retained. Entries whose description matched “frameshift” or “fragment” were excluded.

CharProtDB was downloaded on April 10, 2017. Entries of type trusted_uniprot were excluded as they are redundant with Swiss-Prot. Entries of type trusted_aspgd entries were excluded as they tended to have less specific functional information.

We used MetaCyc release 20.5 and parsed the proteins table (proteins.dat). Only entries with UniProt identifiers and with a reference to at least one paper within the comment field were retained. The corresponding protein sequences were obtained from UniProt. Protein sequences that were fragments were excluded. Protein descriptions corresponding to the MetaCyc entry were obtained from the COMMON-NAME field or from the enzymatic reaction(s) that were linked to each entry. A few entries that lacked names and were not associated with enzymatic reactions were excluded.

REBASE was downloaded on December 8, 2017. REBASE entries were only retained if the sequence specificity of the methyltransferase or nuclease is known. Entries without links to papers were retained because these entries are also usually based on experimental evidence (and not just sequence similarity).

Curated re-annotations of genes’ functions were obtained from the Fitness Browser (http://fit.genomics.lbl.gov/) on November 6, 2018. (These re-annotations are in the Reannotation table in the SQLite3 database, which is available for download (link).) Each entry describes the putative function of a protein-coding gene, as determined from its mutant phenotypes and comparative sequence analysis, along with a rationale. About half of the entries have been described in previous publications (Price, Wetmore, et al. 2018; Price, Zane, et al. 2018).

PaperBLAST also includes another curated database, GeneRIF (Mitchell et al. 2003), which links papers to genes and includes a short summary of the findings about the gene(s). We did not include GeneRIF in Curated BLAST because most of the entries are not descriptions of a protein’s function. For instance, many of the entries are about other aspects of genes such as expression patterns, and many of the entries mention more than one protein. If we had included entries for papers that GeneRIF links to just one protein, then the number of proteins with curated information would increase by 23% (from 115,874 to 149,694). The practical benefit would probably be modest, because 73% of these additional proteins are very similar (over 80% identity and over 75% coverage) to proteins that are curated by other resources.

### Availability of data and source code

The data and the source code for the December 2018 release of Curated BLAST for Genomes and PaperBLAST are available at https://doi.org/10.6084/m9.figshare.7439216.v1. The latest version of the code is available at https://github.com/morgannprice/PaperBLAST.

## Acknowledgements

This material by ENIGMA-Ecosystems and Networks Integrated with Genes and Molecular Assemblies (http://enigma.lbl.gov), a Scientific Focus Area Program at Lawrence Berkeley National Laboratory is based upon work supported by the U.S. Department of Energy, Office of Science, Office of Biological & Environmental Research under contract number DE-AC02-05CH11231

